# Multidirectional alignment of collagen fibers to guide cell orientation in 3D-printed tissue

**DOI:** 10.1101/2025.05.20.654730

**Authors:** Diya Singhal, Fotis Christakopoulos, Lucia G. Brunel, Suraj Borkar, Vanessa M. Doulames, David Myung, Gerald G. Fuller, Sarah C. Heilshorn

**Affiliations:** Department of Chemical Engineering, Stanford University, Stanford, CA, USA; Department of Materials Science and Engineering, Stanford University, Stanford, CA, USA; Department of Ophthalmology, Byers Eye Institute, Stanford University School of Medicine, Palo Alto, CA, USA; VA Palo Alto Healthcare System, Palo Alto, CA, USA

**Keywords:** collagen inks, fiber orientation, extrusion-based 3D printing, cell alignment

## Abstract

Natural tissue comprises fibrous proteins with complex fiber alignment patterns. Here, we develop a reproducible method to fabricate biomimetic scaffolds with patterned fiber alignment along multiple orientations. While extrusion-based approaches are commonly used to align fibrous polymers in a single orientation parallel to the direction of flow, we hypothesized that extrusion-based 3D printing could be utilized to achieve more complex patterns of fiber alignment. Specifically, we show control of lateral spreading of a printed filament can induce fiber alignment that is either parallel or perpendicular to the flow direction. Theoretical prediction of the printing parameters that control fiber orientation was experimentally validated using a collagen biomaterial ink. The velocity ratio of the printhead movement relative to the ink extrusion rate was found to dictate collagen fiber alignment, allowing for the informed fabrication of collagen scaffolds with prescribed patterns of fiber alignment. For example, controlled variation of the ink extrusion rate during a single print resulted in scaffolds with specified regions of both parallel and perpendicular collagen fiber alignment. Human corneal mesenchymal stromal cells seeded onto the printed scaffolds adopted a spread morphology that aligned with the underlying collagen fiber patterns. This technique worked well for filaments either printed into air or extruded within a support bath using embedded 3D printing, enabling the fabrication of 3D structures with aligned collagen fibers. Taken together, this work demonstrates a theoretical and experimental framework to achieve the reproducible fabrication of 3D printed structures with controlled collagen fiber patterns that guide cellular alignment.

## 1. Introduction

Tissues in the body commonly include a fibrous matrix with exquisite fiber alignment along different directions, which guides cell alignment and spreading (1; 2; 3; 4; 5; 6; 7; 8; 9). The most abundant fibrous protein of the native extracellular matrix (ECM) in the human body is collagen, and different, complex patterns of collagen fiber alignment are observed across different tissues, such as the cornea, cartilage, and arteries. (5; 8; 10; 11; 12; 13). For example, in the cornea, the ECM contains orthogonal sheets of collagen fibrils that are able to transmit light while being mechanically resilient (5; 14). In articular cartilage, collagen fibril architecture exhibits depth-dependent organization: fibrils are aligned parallel to the surface in the superficial zone, display a more isotropic distribution in the transitional zone, and orient perpendicular to the surface in the deep zone, where they anchor into the calcified cartilage (12). This zonal arrangement restricts tissue swelling, facilitates osmotic pressurization for load-bearing, and confers both compressive compliance and shear resistance. Arterial collagen fibers are predominantly arranged in a helical pattern, enabling them to reinforce the artery in both the circumferential and axial directions (15; 13).

Collagen fiber alignment can direct many physiological cellular processes, such as migration (16), differentiation (6), and proliferation (17). Thus, a common goal in tissue engineering is to control the orientation of collagen fibers to guide cell behavior (7; 18; 6; 16). To date, several fabrication techniques have been used to create substrates with aligned collagen fibers, including electro-spinning, magnetic flow alignment, microfluidics, mechanical strain devices, and gravity-based fluidic alignment (19; 20; 21; 16; 6). While these aligned collagen constructs have been helpful tools to study cell-matrix interactions such as mechanosensing, cell migration speed, and cell persistence, these techniques are typically limited to unidirectional fiber alignment (22; 16; 23). Creative approaches to overcome this limitation and fabricate radial patterns of collagen fibers have employed Marangoni flow in evaporating, aqueous sessile droplets (24) and microextrusion of discrete filaments in different macroscopic orientations (18). Yet, it remains challenging to achieve larger constructs with controlled fiber alignment in multiple directions.

Recently, 3D printing of biological materials has emerged as a reproducible, scalable, and versatile method to fabricate constructs with clinically relevant dimensions and increased complexity (25; 26; 27; 28; 29; 30; 31; 32). In particular, extrusion-based 3D printing offers ease of use and the potential to produce oriented constructs. Alignment in the extrusion direction has been achieved through different methods for a variety of polymers and fibrous materials. For example, alignment can be induced inside the nozzle by subjecting the material to controlled flow profiles during extrusion; alternatively, alignment can be induced outside of the nozzle by subjecting the printed filament to extensional deformation (33; 34; 7; 35; 36; 37; 38; 39; 40). For fibrous collagen inks, shear stress-induced fiber alignment has been achieved through extrusion-based 3D printing, resulting in constructs with uniaxially aligned fibers in the direction of printing (7; 18).

Here, we sought to develop an easy to implement and reproducible 3D printing strategy to create constructs with collagen fibers aligned in multiple directions. Specifically, we demonstrate the ability to pattern collagen fibers both parallel to and perpendicular to the printing direction, enabling the fabrication of constructs with more complex patterns of fiber alignment. To achieve this, we hypothesized that two different printing regimes could be achieved within a single printing setup. The first regime relies primarily on extensional deformation of the collagen ink within a tapered print nozzle to achieve fiber alignment that is parallel to the print direction. In the second regime, we rationalized that controlling the lateral spreading of the printed filament post-printing can be used to induce collagen fiber alignment perpendicular to the printing direction. To explore this idea, we first theoretically predicted how different printing parameters might impact filament lateral spreading. We then performed a systematic evaluation of the printing parameters for our extrusion-based 3D printer and quantified the collagen fiber alignment patterns for each condition. These experimental observations validated our predictions and allowed us to define the printing parameters for each regime of fiber alignment. We next demonstrated the ability of these parallel and perpendicularly aligned collagen substrates to guide the spreading morphology of cells. Using this new printing strategy, we fabricated specimens with multidirectional collagen fiber alignment in a single print. Taken together, this work introduces a predictable, reproducible, and scalable approach to print biomimetic constructs with complex patterns of collagen fiber directionality to guide cellular alignment.

## 2. Results and discussion

### 2.1. Collagen fiber alignment along the printing direction

Collagen inks are widely used in 3D printing, both in the solution and the gel state (10; 41; 42; 7). Collagen gelation is attained through physical assembly and entanglement of collagen fibers at physiological conditions, with the kinetics being dependent on collagen concentration, temperature, and pH (43; 44; 45; 46). Collagen inks have been formulated to form fibers before printing, during printing, or after printing (10).

We chose to formulate a collagen ink with pre-formed fibers that would become oriented by fluid mechanics, as we reasoned that printing parameters would have a stronger impact on pre-formed fibers as opposed to soluble collagen protein (47). Specifically, we selected a neutralized type-I atelocollagen at a relatively high concentration of 35 mg mL^−1^ that is stored and printed at 4^°^C. At these conditions, the ink has storage (*G*^′^) and loss (*G*^′′^) moduli of about 250 Pa and 150 Pa, respectively (figure 1(A)). Thus, the ink is already a weak gel (*G*^′^ > *G*^′′^) prior to printing, due to its high concentration and neutral pH. Importantly, the ink at 4^°^C displays shear-thinning behavior, making it suitable for extrusion-based 3D printing (figure 1(B)). Additionally, this ink formulation demonstrates the ability to recover after shear-thinning. After exposure to high shear that disrupts the network structure, the fluid achieves a 80% storage modulus recovery in 30 seconds at 4^°^C and recovers its initial stiffness after 240 seconds(supplemental figure S1(A)). As the ink is deposited on the printbed, which is at room temperature, the physical assembly of the collagen fibers continues to progress, as evident in the ten-fold increase in *G*^′^, reaching a plateau of 2,000 Pa (figure 1(C)). The presence of fibers in the ink cartridge was confirmed by confocal reflectance microscopy at 4^°^C (figure 1(D)). As expected, collagen fibers within the ink are initially randomly oriented.

**Figure 1.**
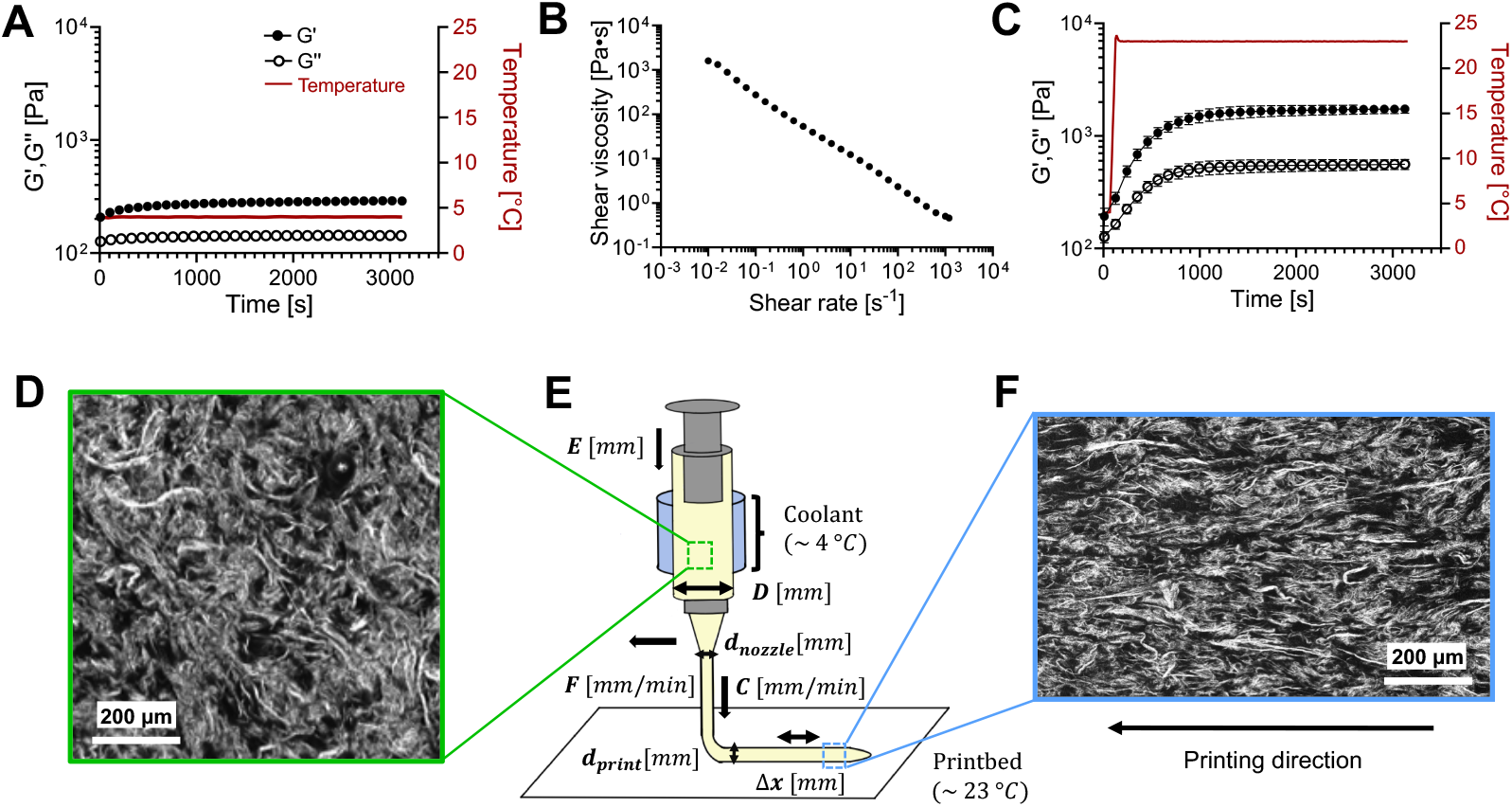
Extrusion-based 3D printing setup for collagen fiber alignment. (A) Time sweep, shear rheology of the collagen ink at 4^°^C; (storage modulus, *G*^′^ , •; loss modulus, *G*^′′^, °; temperature: red solid line). (B) Shear-dependent viscosity of the collagen ink at 4^°^C. (C) Shear rheology of the collagen ink as temperature increases from 4 to 23^°^C; (storage modulus, *G*^′^ , •; loss modulus, *G*^′′^, °; temperature: red solid line). (A-C) Data are mean ± standard deviation; N = 3 independent gel conditions. For (A) and (B), symbol size is larger than the errorbars. (D) Representative confocal reflectance image of the collagen ink prior to extrusion. (E) Schematic of the extrusion-based printing setup showing the different printing parameters; ink cartridge diameter (*D*, mm), nozzle diameter (*d*_nozzle_, mm), translational speed of the printhead (*F*, mm min^−1^), speed of ink extrusion (*C*, mm min^−1^), plunger displacement per step (*E*, mm), lateral displacement distance of the printhead per step (Δ*x*, mm), and diameter of the printed filament (*d*_print_). An ice pack is positioned around the syringe to keep the ink at 4^°^C, which is deposited on a printbed in air at room temperature. (F) Representative confocal reflectance image of the collagen post-printing, showing fiber alignment parallel to the printing direction (black arrow).

To demonstrate that this collagen ink could achieve parallel alignment similar to that reported by others (7), we employed a standard extrusion-based 3D printing system (figure 1(E)). This setup offers several user-defined parameters including the ink cartridge diameter, *D*, which we kept constant through all experiments (7.29 mm), and the diameter of the printing nozzle, *d*_nozzle_, which was varied in our studies. In addition to these hardware parameters, the setup also includes a number of user-defined parameters controlled through the software (*e*.*g*. Gcode). These include the plunger displacement per step, *E*, the lateral displacement of the printhead per step, Δ*x*, and the translation speed of the printhead, *F*. Through these hardware- and software-controlled variables, we define further printing parameters such as the speed of ink extrusion from the nozzle, *C*, and the diameter of the printed filament, *d*_print_. Through optimization, we were able to identify a combination of printing parameters (supplemental table S1) that results in parallel fiber alignment at within the printed filament at 23^°^C (figure 1(F)), similar to results of others (7). Upon further heating of the collagen print to physiological temperature (37^°^C), additional collagen fiber formation occurs, as evidenced by a further increase in *G*^′^ to about 3500 Pa (supplemental figure S1(B,C)). To confirm that the heating to 37^°^C does not disrupt the achieved fiber alignment, we incubated the sample for 15 minutes and then confirmed that the alignment remains unchanged by confocal reflectance imaging (supplemental figure S1(D)). Having been able to replicate the state of the art with our extrusion-based 3D printing setup, we next investigated the ability to achieve fiber alignment in an orientation other than the printing direction, towards the broader goal of fabricating constructs with multidirectional collagen fiber alignment.

### 2.2. Printed filament spreading dictates fiber alignment orientation

Previously, an interesting correlation between extruded width and fiber alignment has been shown where low extrusion width lead to more fibers aligned in printing direction however as extruded width increases less fibers are aligned in the printing direction (48). We hypothesized that harnessing the lateral spreading of the printed filament post-printing would lead to perpendicular alignment of the collagen fibers. This lateral spreading of the printed filament can be quantified as a ratio of the printed filament diameter (*d*_print_) to the nozzle diameter (*d*_nozzle_), which we define as the diameter ratio, *D*^*^, (supplemental figure S2):

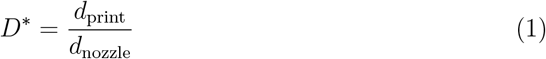

Thus, when *D*^*^ *<* 1, the printed filament diameter is smaller than the nozzle diameter, implying less printed filament spreading. In contrast, when *D*^*^ *>* 1, the printed filament diameter is higher than the nozzle diameter, denoting more printed filament spreading. To control the lateral spreading of the printed filament, we derived a theoretical expression of *D*^*^ by performing a mass balance. For our extrusion-based, piston-controlled, 3D printer, the volumetric flow rate inside the syringe 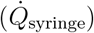 is defined as:

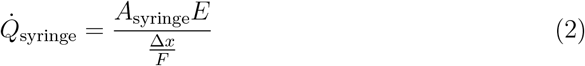

which, due to mass conservation, is equal to the volumetric flow rate at the end of the nozzle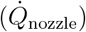:

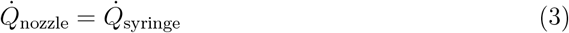

Similarly, the volumetric flow rate at the end of the nozzle can be equated to that of the printed filament after extrusion from the nozzle:

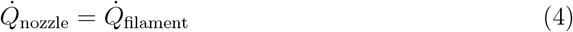

We initially assume the printed filament is a cylinder to enable the estimation of 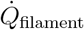 as:

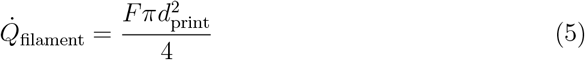

In actuality, the printed filament will not be a perfect cylinder, and to account for this, we introduce the coefficient, *β*. When *β* = 1, then the cross-sectional area of the printed filament would be perfectly circular (*i*.*e*. the width and height of the printed filament will be similar), and when *β >* 1 , the filament will undergo lateral spreading.

Using conservation of mass, through equations 2-5 we find:

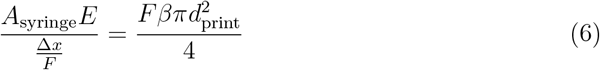

giving us a theoretical prediction for *d*_print_ as:

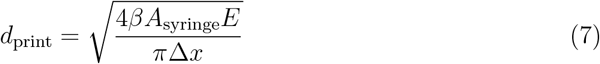

In turn, *D*^*^ (*i*.*e*. the normalized printed filament diameter) can also be correlated to the printing parameters as:

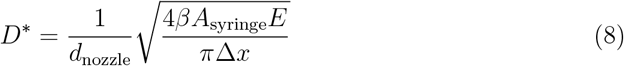

From equation 8, it can be duly noted that *D*^*^ depends on two of the printing parameters introduced in figure 1(E), namely *E* (plunger distance per step) and *d*_nozzle_ (nozzle diameter), with the proportionality given as:

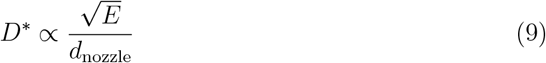

To test this theoretical prediction, we conducted a series of controlled experiments wherein either *E* or *d*_nozzle_ was independently varied. For each experimental condition, we measured the actual *d*_print_ (*i*.*e*. the printed filament width) through confocal imaging to calculate an experimentally observed *D*^*^. First, we kept *E* constant (*E* = 0.007 mm) and decreased *d*_*nozzle*_ from 1.55 mm to 0.16 mm (a ratio of ≃ 9.7:1). Upon microscopy analysis of the resulting printed filaments, we observed that decreasing *d*_nozzle_ causes *D*^*^ to increase from 0.875 to 8.406 (a ratio of ≃ 1:9.6) (figure 2(A)). Thus, for these conditions, we found that *D*^*^ and *d*_*nozzle*_ are inversely proportional, consistent with our theoretical prediction (equation 9).

**Figure 2.**
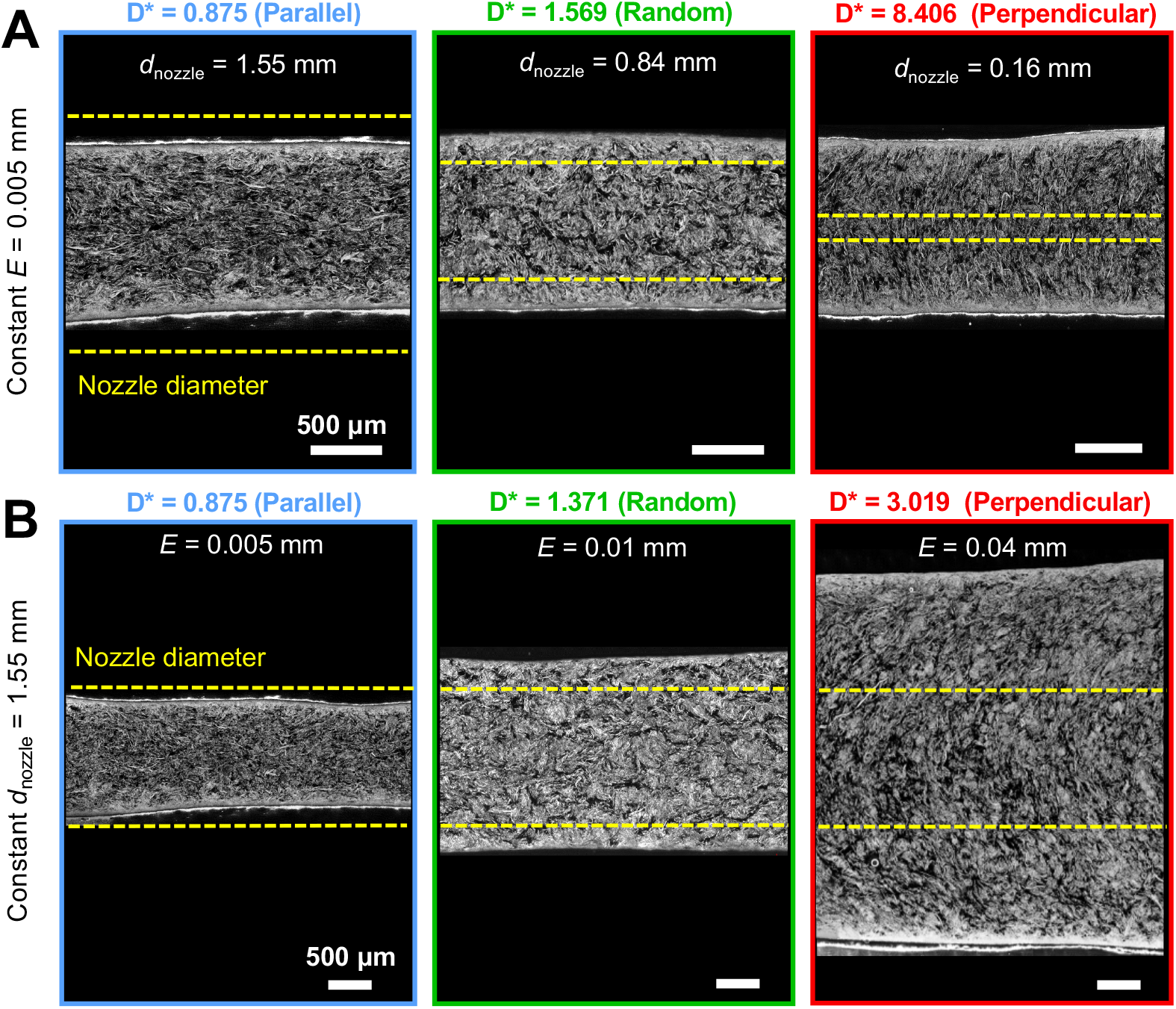
Diameter ratio correlates with collagen fiber alignment in printed filaments. (A) Representative confocal reflectance images of collagen filaments printed with the same plunger displacement per step (*E* = 0.005 mm) and different nozzle diameters (*d*_nozzle_). (B) Collagen filaments printed with the same nozzle (*d*_nozzle_ = 1.55 mm) and different plunger displacement per step (*E*). Yellow dotted lines denote the nozzle diameter for easy comparison to the printed filament width. The experimentally determined diameter ratio, *D*^*^, (*i*.*e*. the width of the printed filament, *d*_print_, normalized by the diameter of the printing nozzle, *d*_nozzle_) is written above each micrograph along with a qualitative description of the observed collagen fiber orientation. Scalebars correspond to 500 *µ*m.

Next, we kept *d*_nozzle_ constant (*d*_nozzle_ = 1.55 mm) while *E* was increased from 0.005 mm to 0.040 mm (a ratio of ≃ 1:8) (figure 2(B)). Upon measuring the width of the printed filament for increasing *E*, we observed that *D*^*^ increased from 0.875 to 3.019 (a ratio of ≃ 1:3.45). Thus, these experimental results are consistent with our theoretical prediction that *D*^*^ is directly proportional to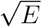 (equation 9).

After validating our ability to predict the printed filament width based on our printing parameters, we then observed the fiber alignment within each filament using confocal reflectance microscopy. To aid in the assessment of fiber alignment direction and to prevent observational bias, we used an automated fiber tracing algorithm (CT-FIRE for Individual Fiber Extraction) that traces and artificially colors individual fibers to enable visualization of fiber orientation (supplemental figure S3). We found that at low *D*^*^ (*i*.*e*. the condition where uniaxial filament extension occurs, 0.875 in the left panels in figure 2(A,B)), the collagen fibers were oriented parallel to the printing direction (supplemental figure S3).

In contrast, for the conditions with *D*^*^ *>* 1 (*i*.*e*. significant lateral filament spreading, 8.406 and 3.019 in the right panels of figure 2(A) and 2(B), respectively) the collagen fibers were observed to orient perpendicular to the filament printing direction (supplemental figure S3). This observation is consistent with our hypothesis that fluid flow that occurs after extrusion while the filament is undergoing uniaxial extension or lateral spreading can be sufficient to induce parallel or perpendicular alignment of embedded collagen fibers, respectively. For experimental conditions with an intermediate value of *D*^*^ (*D*^*^ ≃ 1, middle panels of figure 2(A,B)), the collagen fibers were randomly oriented (supplemental figure S3). This observation suggests that there is an intermediate transition regime where the embedded fibers experience neither sufficient extensional flow to orient in the parallel direction, nor enough lateral flow to orient perpendicularly.

### 2.3. Quantification of fiber alignment across a broad printing space

We next wanted to identify the relevant values of *D*^*^ to achieve parallel or perpendicular collagen fiber alignment across a wide range of printing parameter space. To categorize the prints into different alignment regimes, we developed a technique to quantify the fiber alignment orientation using automated software analysis from the confocal reflectance micrographs (figure 3(A)). From this analysis, we prepared histograms showing the normalized frequency of fiber alignment relative to the total number of fibers versus fiber orientation, where the printing direction is 0^°^ (figure 3(B, C) and supplemental figures S4(B), S5(A,B)).

**Figure 3.**
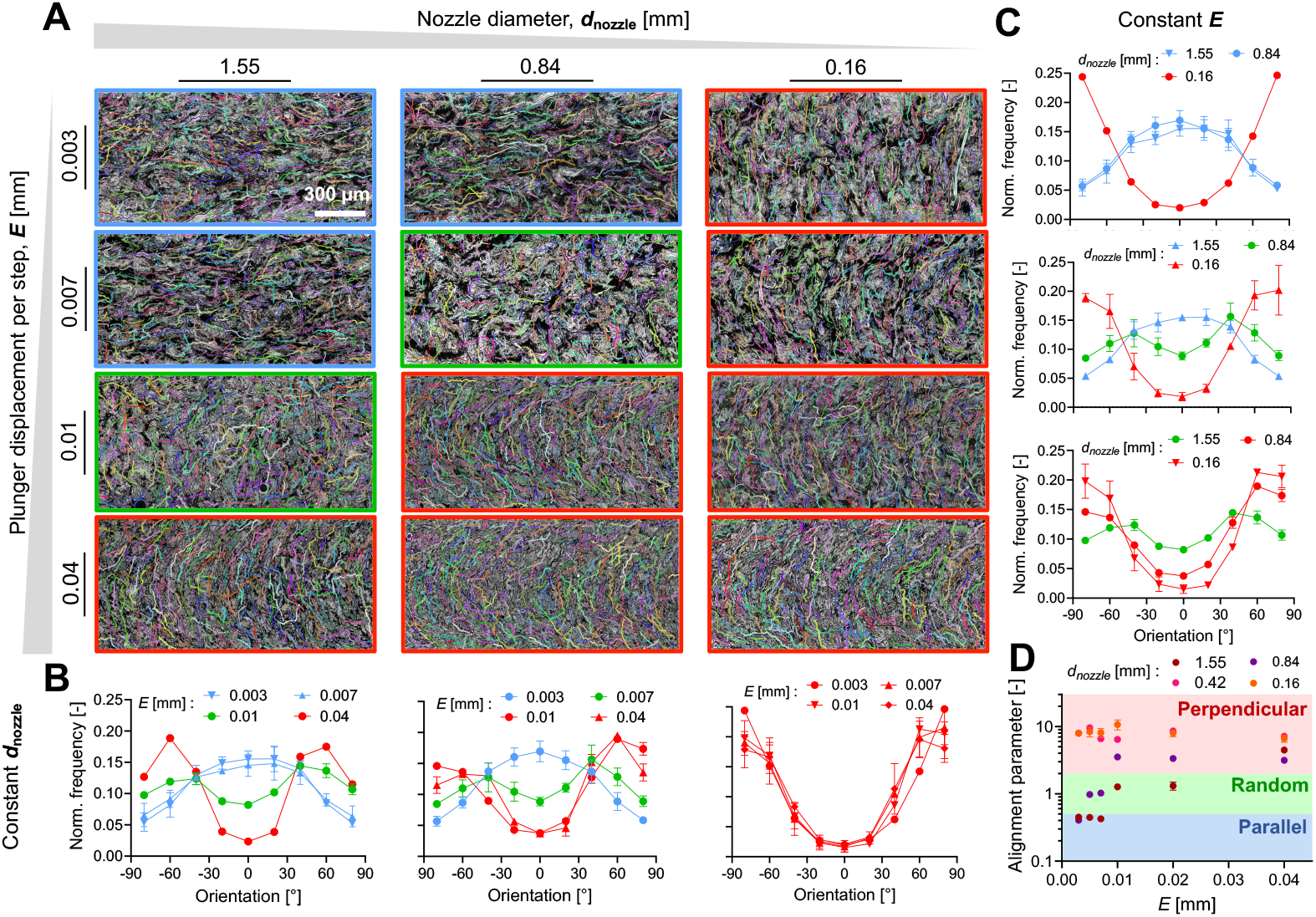
Selection of printing parameters controls the collagen fiber alignment direction. (A) Automated analysis of representative confocal reflectance images for different combinations of plunger displacement per step (*E*) and nozzle diameter (*d*_nozzle_) leads to identification of parallel (blue), random (green) and perpendicular (red) alignment regimes. (B) Histograms of fiber alignment when keeping the plunger displacement constant and changing the nozzle diameter, showing the normalized frequency of fiber alignment relative to the total number of fibers versus fiber orientation, where the printing direction is 0^°^. (C) Histograms of fiber alignment when keeping the nozzle diameter constant and changing the plunger displacement. (D) Alignment parameter (*AP*) plotted for several combinations of *E* and *d*_nozzle_ show that the printing parameter space can be divided into three regimes with different fiber alignment directions: parallel (blue, *AP* ≤ 0.5), random (green, 0.5 *< AP* ≤ 2) and perpendicular (red, *AP >* 2). (B,C) Data are mean ± standard deviation, N = 3 independent printing trials.

From these histograms, we further calculated an Alignment Parameter (*AP*), which we define as the ratio of the fraction of perpendicular fibers (orientation ranges from −80^°^ to −60^°^ and 60^°^ to 80^°^) to the fraction of parallel fibers (orientation from −20^°^ to 20^°^). To support visual validation of our automated analysis, we also used the aforementioned automated fiber tracing algorithm to artificially color individual fibers (figure 3(A)). With this analytical workflow in place, we systematically varied *E* and *d*_nozzle_ and quantified the fiber alignment for three independent printing trials at each of 24 conditions. First, we found that by keeping *d*_nozzle_ constant ( *d*_nozzle_ = 1.55 mm or 0.84 mm, left and middle columns, respectively, in figure 3(A)), the fiber alignment direction can be transitioned from parallel to random to perpendicular by increasing the plunger displacement *E* from 0.003 mm to 0.04 mm (left and middle graph in figure 3(A,B) and supplemental figure S4). Interestingly, for choice of smaller *d*_nozzle_ (0.16 mm, right-most column, (figure 3(A)), the fiber alignment direction was predominantly perpendicular for all *E* values tested (right column in figure 3(A,B)). These data suggest that careful selection of a single nozzle could be used to print all three different fiber orientations within a single construct.

An alternative method to control the fiber alignment orientation is to keep *E* constant while varying *d*_nozzle_. For lower *E* values (0.003 or 0.007 mm, top two rows in figure 3(A)), decreasing *d*_nozzle_ from 1.55 to 0.16 mm resulted in a transition from parallel to perpendicular fiber alignment, respectively (top two graphs in figure 3(C) and supplemental figure S5(A)). For larger *E* (0.01 mm or 0.04 mm, bottom two rows in figure 3(A)), the alignment direction is predominantly perpendicular for all *d*_nozzle_ values (bottom graph in figure 3(C)) and supplemental figure S5(B)). These data suggest that multinozzle printing systems also could be used to achieve all three different fiber orientations within a single construct.

To synthesize together the information from each histogram into a single quantitative metric, we next calculated and plotted the *AP* value for each of the 24 tested combinations of printing parameters (figure 3(D)). This allowed us to identify cutoff values of *AP* that define the three regions of fiber alignment: parallel (blue, *AP* ≤ 0.5), random (green, 0.5 < *AP* ≤ 2) and perpendicular (red, *AP* > 2) (figure 3(D)). At the cutoff value of AP ≤ 0.5 , the number of fibers aligned in the parallel direction is approximately twice that of the perpendicular direction. In the green, random region (0.5 < AP < 2), the proportions of parallel and perpendicular fibers are comparable. Finally, for AP ≥ 2, roughly twice as many fibers are oriented perpendicularly compared to those aligned in the parallel direction. Interestingly, for this particular ink formulation and printer setup, we found that 16 of the 24 tested printing parameter combinations resulted in perpendicular fiber alignment (figure 3(D)). Thus, while most bioprinting studies to date have focused on the parallel alignment of fibers, *printing conditions to achieve perpendicular fiber alignment are readily achievable through our printing strategy*.

## 2.4. Normalized speed of ink extrusion dictates fiber alignment orientation

Although the exact values of *E* and *d*_nozzle_ that result in perpendicular or parallel fibers will be different for each printer setup, we reasoned that more generalizable guidelines could be identified by considering how *AP* relates to the *D*^*^ ratio as defined previously in equation 8. Towards this goal, we first experimentally identified the dependence of *E* and *d*_nozzle_ on the printed filament width (*d*_print_). As *E* increases from 0.003 to 0.04 mm, *d*_print_ increases monotonically due to the increased amount of extruded material. Interestingly, we observed that for a particular *E* value, *d*_print_ remains the same irrespective of *d*_nozzle_ (figure 4(A)), which is consistent with our theoretical prediction (equation 7). By plotting *AP* against *D*^*^ for our 24 printing conditions, we were able to identify the ranges of *D*^*^ for the three fiber alignment regimes using our earlier *AP* cutoff values (figure 4(B)). Specifically, we found that *D*^*^ ≤ 1 produced filaments in the parallel fiber alignment region (*AP* ≤ 0.5), and *D*^*^ *>* 2 produced filaments in the perpendicular fiber alignment region (AP ≥ 2). For values between these two regions, *i*.*e*. 1 *< D*^*^ ≤ 2, fibers had random alignment.

**Figure 4.**
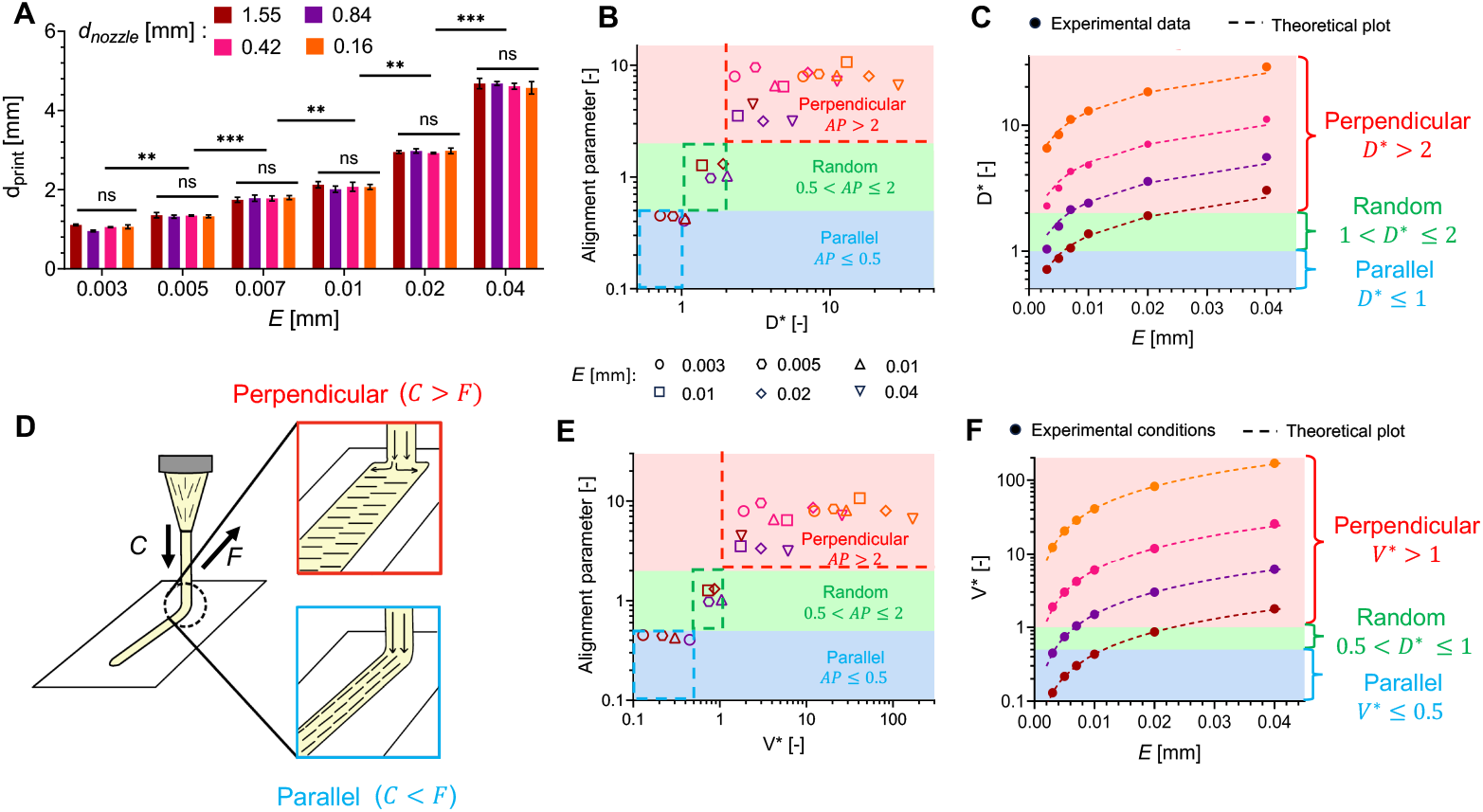
Relative speed of ink deposition controls filament spreading and fiber alignment. (A) Experimentally determined printed filament width (*d*_print_) plotted as a function of *E* for different values of *d*_nozzle_. (B) Alignment parameter (*AP*) plotted against *D*^*^ across 24 printing conditions, with distinct alignment regimes based on *AP*. Symbol colors denote *d*_nozzle_, while symbol shapes denote *E*. (C) Experimental data and theoretical fits for *D*^*^ as a function of *E* for different values of *d*_nozzle_. (D) Schematic of fiber alignment regimes for different velocity ratios (*V*^*^), where *V*^*^ is the ratio of ink extrusion speed from the nozzle (*C*) to the translational speed of the printbed (*F*). (E) Alignment parameter (*AP*) plotted against *V*^*^ for 24 printing conditions, with distinct alignment regimes based on *AP*. (F) Theoretical values of *V*^*^ plotted as a function of *E* for different values of *d*_nozzle_. (A,B,C,E,F) Data are mean ± standard deviation, N = 3 independent printing trials.^**^*p <* 0.01,^***^ *p <* 0.001,^****^ *p <* 0.0001.

We then compared the experimentally determined values of *D*^*^ to our earlier theoretical prediction (equation 8), where the only fitting parameter is *β* (figure 4(C)). When fitting our experimental data to this equation, we found the same value of *β* = 4 for each *d*_nozzle_ tested across the full range of *E*. This constant fitting parameter further supports the notion that *β* is an ink-dependent parameter that does not change with the tested printing conditions, and thus remains unchanged for a given ink system (*e*.*g*. ink composition and temperature).

To reformulate our theoretical prediction into a format that is independent of *β*, so that it is controlled solely by the experimental printing parameters (*e*.*g. E, d*_nozzle_, *F* , *D*, Δ*x*) for a specific ink system, we defined the normalized velocity, *V*^*^:

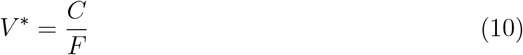

where *C* corresponds to the speed of the ink extrusion from the nozzle and *F* to the translational speed of the printhead. We hypothesized that when *C* >> *F* , *i*.*e*. the speed at which ink is extruded is higher than the translational speed of the printhead, the ink would experience significant lateral spreading on the printbed, leading to perpendicular fiber alignment relative to the printing direction (figure 4(D)). In contrast, when *C* << *F* , *i*.*e*. the speed at which the ink is extruded is lower than the translational speed of the printhead, the filament will become extensionally drawn as it gets extruded, leading to parallel fiber alignment. When *C* ≤ *F* , *i*.*e*. the speed at which ink is extruded is comparable to the translational speed of the printhead, the filament does not experience significant lateral spreading or extensional drawing, leading to random alignment of fibers.

This normalized velocity framework enables extension to other extrusion-based systems such as pneumatic-controlled printers, where *F* is explicitly selected, while *C* will depend on the pressure drop through the syringe and nozzle, Δ*P*. The maximum stress, *τ*_*w*_ in capillary flow can be calculated from: (49)

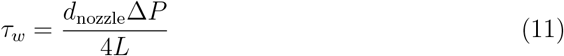

where *L* is the length of the nozzle. For power-law fluids, the shear rate, 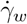, calculated from:

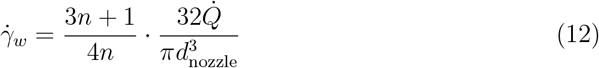

where *n* is the power-law index. Through supplemental equation (S4) and:

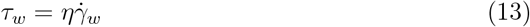

where *η* is the apparent shear viscosity, we can get an expression for *C*:

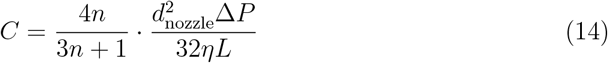

For our specific extrusion-based 3D printer, the theoretical equation for *V*^*^ is derived using mass conservation by equating the volumetric flow rate inside the syringe to the volumetric flow rate at the tip of the printing nozzle (supplemental equations S1-S6) and can be expressed as:

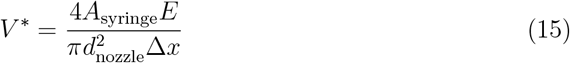

To confirm that this formulation of *V*^*^ can directly predict the direction of fiber alignment without any fitting parameters, we correlated the resulting theoretical *V*^*^ values with the experimentally determined alignment parameter (*AP*) across the range of *E* and *d*_nozzle_ conditions presented earlier (figure 4(E)). As before, we used our *AP* cutoff values to identify the regimes that correspond to perpendicular, random, and parallel fiber alignment. From this analysis, we observed that a *V*^*^ *<* 0.5 results in parallel fibers (*AP* ≤ 0.5), intermediate values of *V*^*^ (0.5 ≤ *V*^*^ ≤ 1) results in randomly aligned fibers (0.5 *< AP* ≤ 2), and *V*^*^ ≥ 1, results in perpendicular fibers (*AP >* 2). These results agree with a similar ratio that was identified in earlier studies for an extrusion deposition additive manufacturing system (the ratio of the extrusion speed and print speed), where ratios less than unity resulted in greater fiber alignment in the printing direction (50).

To enable the prediction of fiber alignment with no fitting parameters, we plotted the theoretical curves of *V*^*^ as a function of *E* for each *d*_nozzle_ tested, using equation 15 (figure 4(F)). For ease of viewing, the three different fiber alignment regimes are shown as different color-coded regions, and we overlaid the points corresponding to the 24 specific combinations of *E* and *d*_*nozzle*_ tested out experimentally. Similar plots could be constructed for other printer-ink combinations to enable prediction of printing parameters that result in specific fiber alignment orientations. As extrusion-based printers are generally widely available and easy to use, this framework could be readily adapted to other systems. While the diameter ratio (*D*^*^) included a fitting parameter that is related to the intrinsic ink properties, the velocity ratio (*V*^*^) depends only on the printing parameters programmed into the 3D printer.

### 2.5. Control of fiber alignment in embedded 3D printing

As our data suggest that fiber alignment is primarily dictated by filament spreading versus filament drawing (and does not depend on interactions between the filament and the printbed), we hypothesized that the same printing parameters that induce parallel or perpendicular fiber alignment when printed in air would also cause fiber alignment when printed in a support bath. Embedded 3D printing into support baths has emerged as a strategy to expand the range of printable materials and the complexity of fabricated structures with intricate features (51; 52; 53). In contrast to printing in air, in embedded 3D printing the ink material is deposited into an aquaeus yield-stress fluid (51). This allows the printed material to remain suspended and supported during the printing process, enabling the creation of complex 3D structures that would otherwise collapse under their own weight and/or aqueous surface tension if printed in air (51).

To explore if our fiber alignment strategy could be extended to embedded 3D printing, we prepared a Carbopol support bath material. Carbopol is widely used as a support bath due to its shelf-healing behavior (figure 5(A)) (54). In addition, previous work has demonstrated that a large range of printing speeds are compatible with Carbopol support baths without causing porosity in the final print (55).

**Figure 5.**
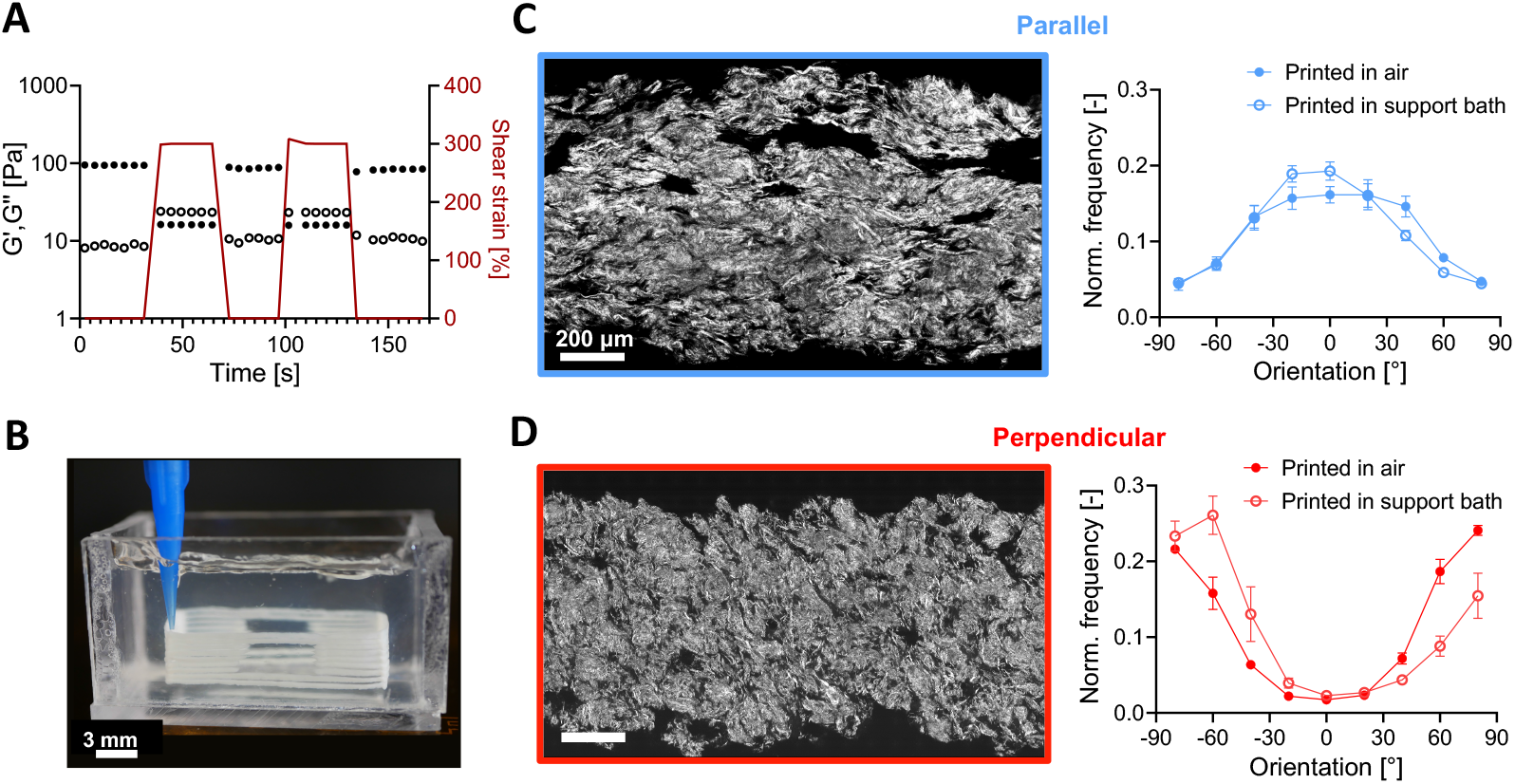
Collagen fiber alignment in embedded 3D printing. (A) Shear rheology of Carbopol support bath upon alternating application of high or low shear strain (red line) demonstrates shear-thinning and shelf-healing behavior. Filled symbols correspond to storage moduli, *G*^′^, and open symbols correspond to loss moduli, *G*^′′^. (B) Multilayer collagen construct 3D printed in a Carbopol support bath. (C-D) (Left) Representative confocal reflectance images with parallel (C) and perpendicular (D) fiber alignment. (Right) Quantification of the collagen fiber alignment of collagen filaments printed in a Carbopol support bath compared to controls printed in air at the same conditions. (C,D) Data are mean ± standard deviation; n=3 independent printing experiments.

As a demonstration, we printed a 3D structure having hollow, suspended features (*i*.*e. windows*) and built up as a multilayered construct in which fiber alignment was tuned in each individual filament (figure 5(B)). In addition, single filaments, which allow for easier visualization of the collagen fibers after removal from the support bath, were printed with identical printing parameters (supplemental table S1). The fiber alignment of the resulting prints was imaged and quantified for comparison to that of filaments printed in air with the same printing conditions (figure 5(C,D)). Collagen fibers were well aligned when filaments were printed into the Carbopol support bath, both for conditions that result in parallel and perpendicular orientations. Furthermore, the degree of fiber alignment was found to be similar for filaments printed in air or within the Carbopol support bath. For example, the *AP* for parallel alignment printing in air and in the support bath was found to be 0.35 ± 0.044 and 0.30 ± 0.02, respectively. The *AP* for perpendicular alignment in air and in the support bath was found to be 6.23 ± 0.75 and 9.57 ± 0.017, respectively. While printing in a support bath can produce filaments with both parallel and perpendicular fiber alignment, the inherent limitations and challenges of embedded 3D printing must also be considered when selecting printing parameters. For example, moving the printhead quickly (*V*^*^ *<* 0.5) can result in parallel fiber alignment, but this speed may also introduce print defects such as filament breakup, “crowning” (*i*.*e*. upwards flow of the ink material due to a prolonged crevice in the support bath), and extensive porosity (51; 55; 56; 57). Therefore, the printing parameter regimes for each alignment orientation will be affected by the viscosity ratio between the ink material and the support bath, which will need to be tuned for each material system.

### 2.6. Collagen fiber alignment guides cell orientation

To evaluate if patterned collagen fibers could influence cellular orientation, we seeded human corneal mesenchymal stromal cells (CMSCs) on collagen scaffolds with parallel, perpendicular, and random fiber alignments. The native cornea has multidirectional collagen fiber alignment, which is responsible for imparting its structural and functional properties (58; 59; 5; 60), and CMSCs are one of the regenerative cell types found in these tissues (61; 62; 63). Specimens were printed with parallel or perpendicular collagen fiber alignment, following the printing guidelines identified earlier (parallel: *V*^*^= 0.22, *d*_nozzle_= 1.55 mm; perpendicular: *V*^*^= 2.98, *d*_nozzle_= 0.42 mm), and a cast gel condition with random fiber alignment was included as a negative control. The collagen fibers in the three different conditions were imaged through confocal reflectance on day 0, prior to cell seeding (first column of figure 6(A) and supplemental figure S6), and the fiber orientation was determined for each sample. At days 1, 3, and 5 after CMSC seeding, the cell morphology was visualized with a live cell cytoplasmic marker (calcein-AM). Across all timepoints, cells remained highly viable (about 99%, supplemental figure S7) and maintained their proliferative phenotype, resulting in increased confluence by day 5. Cell alignment was quantified by calculating orientation histograms from these images on days 1 and 3. Already by day 1, quantified cell orientation matched with the underlying collagen fiber alignment across all three types of fiber patterns (figure 6(B-D)). This agreement in alignment orientation persisted at day 3, as quantitatively evaluated in figure 6(B-D). At day 5, increased cell-cell contact due to cell proliferation prevented automated identification of individual cell borders for orientation analysis; nonetheless, fiber-guided cell alignment was readily observed through qualitative inspection of the confocal fluorescence micrographs. Cell morphology was further analyzed by staining the cell nuclei (DAPI) and filamentous actin (phalloidin) along with the aldehyde dehydrogenase 3 family member A1 (ALDH-3A1), a phenotypic marker for viable CMSCs (figure 6(A)) (64). Across all three cases (perpendicular, parallel, and random fiber alignments), and all timepoints, cells were observed to elongate and align with the underlying substrate while maintaining their positive staining for ALDH-3A1. Thus, similar to reports by others, controlling the orientation of collagen fibers at the microscale allows control of the cell orientation (7; 22; 6), which we uniquely demonstrate here by inducing cells to align in both parallel and perpendicular orientations.

**Figure 6.**
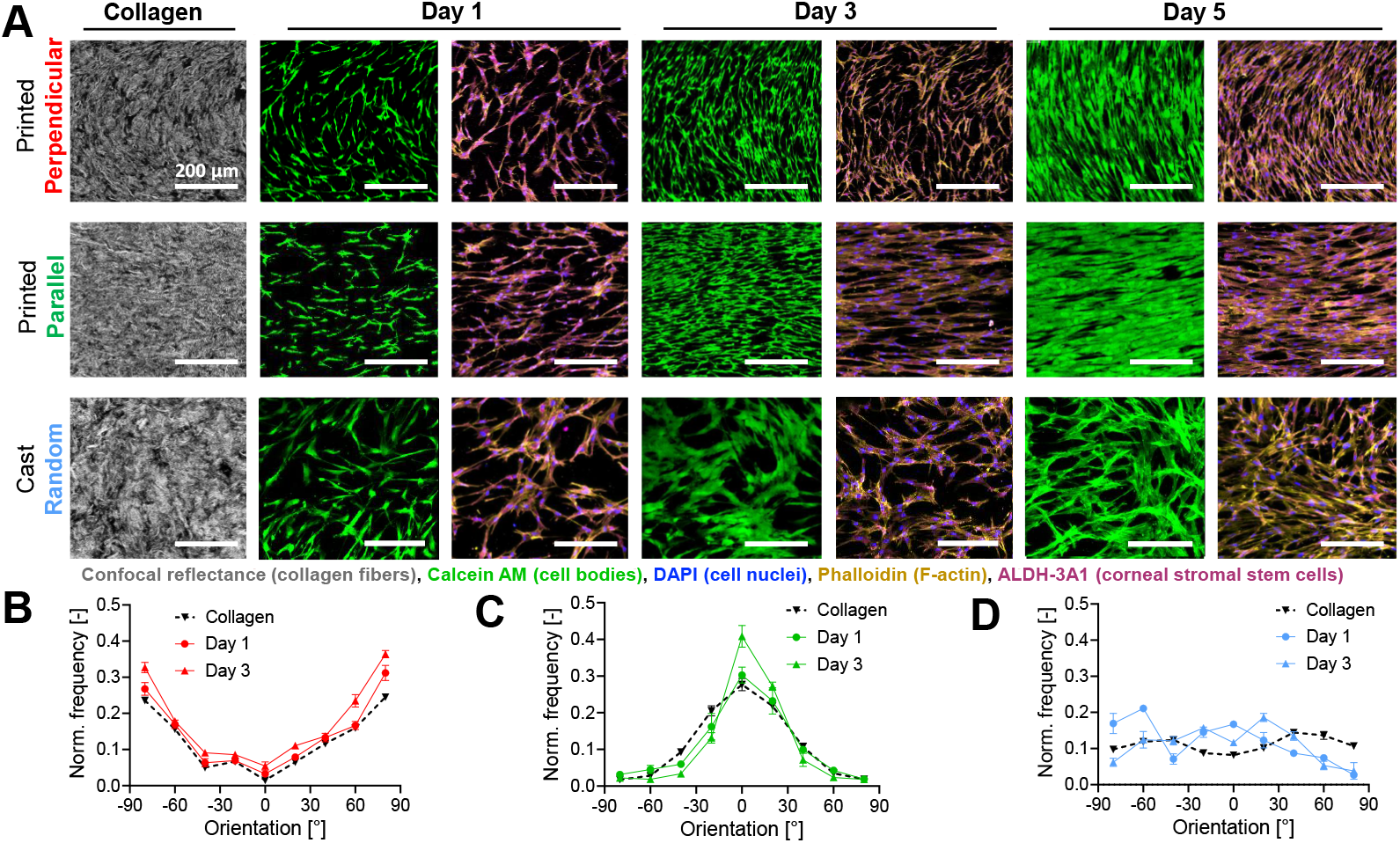
Collagen fiber alignment guides cell orientation. (A) Representative confocal reflectance images of collagen fibers (gray) and confocal fluorescence images of CMSCs grown on the collagen filaments for 5 days and stained for cell bodies (green, calcein-AM), cell nuclei (blue, DAPI), filamentous actin (orange, phalloidin), and ALDH-3A1 (magenta). Printed samples included both perpendicular (top row) and parallel (middle row) collagen fiber alignment, and controls included cast samples (bottom row) with random fiber alignment. (B-D) Orientation of the the collagen fibers and seeded cells on days 1 and 3 were quantified for the prints with perpendicular (B), parallel (C), and random (D) fiber alignments, normalized to the total count of fibers or cells, respectively.

### 2.7. Multidirectional collagen fiber alignment in a single print

In native tissues, collagen fibers and endogenous cells are aligned in multiple directions within a single tissue (5; 8; 10; 11; 12; 13). Inspired by this, we reasoned that we should be able to modulate the printing conditions (*i*.*e. V*^*^) during the fabrication process to achieve multidirectional collagen fiber alignment and hence different regions of cellular orientation within a single construct. Specimens with three different designs were created to demonstrate multidirectional alignment through either different interfilament or intrafilament fiber orientation patterning (figure 7(A)). In one type of interfilament patterning, sequential filaments can have alternating parallel and perpendicular fiber alignment. This can be achieved either (i) by using the same *E* but printing through two differently sized nozzles (left column, figure 7(A) and supplemental table S1) or (ii) by printing through the same nozzle (*d*_*nozzle*_ = constant) and varying *E* for sequential filaments (middle column, figure 7(A) and supplemental table S1). In intrafilament patterning, alternating regions of parallel and perpendicular fiber alignment are present in the same filament. This can be achieved using a single nozzle by varying *E* during continuous printing of a filament (right column, figure 7(A) and supplemental table S1). CMSCs were seeded onto these three different multidirectional fiber alignment prints, and cell morphology was visualized through cytoplasmic calcein staining on day Cellular orientation was digitally analyzed and color coded to identify cells with parallel (blue, -45^°^ to 45^°^,) or perpendicular alignment (red, -90^°^ to -45^°^ and 45^°^ to 90^°^) relative to the printing direction (figure 7(B)). Cells appeared to adhere equally well on construct regions with parallel or perpendicular fiber alignment, (figure7(C)). As expected, cells followed the alignment of the underlying collagen fibers, as visualized by automated color coding (figure 7(D)) and by quantitative analysis (figure 7(E)). Thus, this technique offers the flexibility to create multidirectional patterns of fiber alignment using either a multiple nozzle printer (by altering *d*_nozzle_ and keeping *E* constant) or a single nozzle printer (by altering *E* and keeping *d*_nozzle_ constant). This versatile approach will enable the fabrication of printed constructs with control over different regions of patterned cell orientation to mimic native tissue architecture.

**Figure 7.**
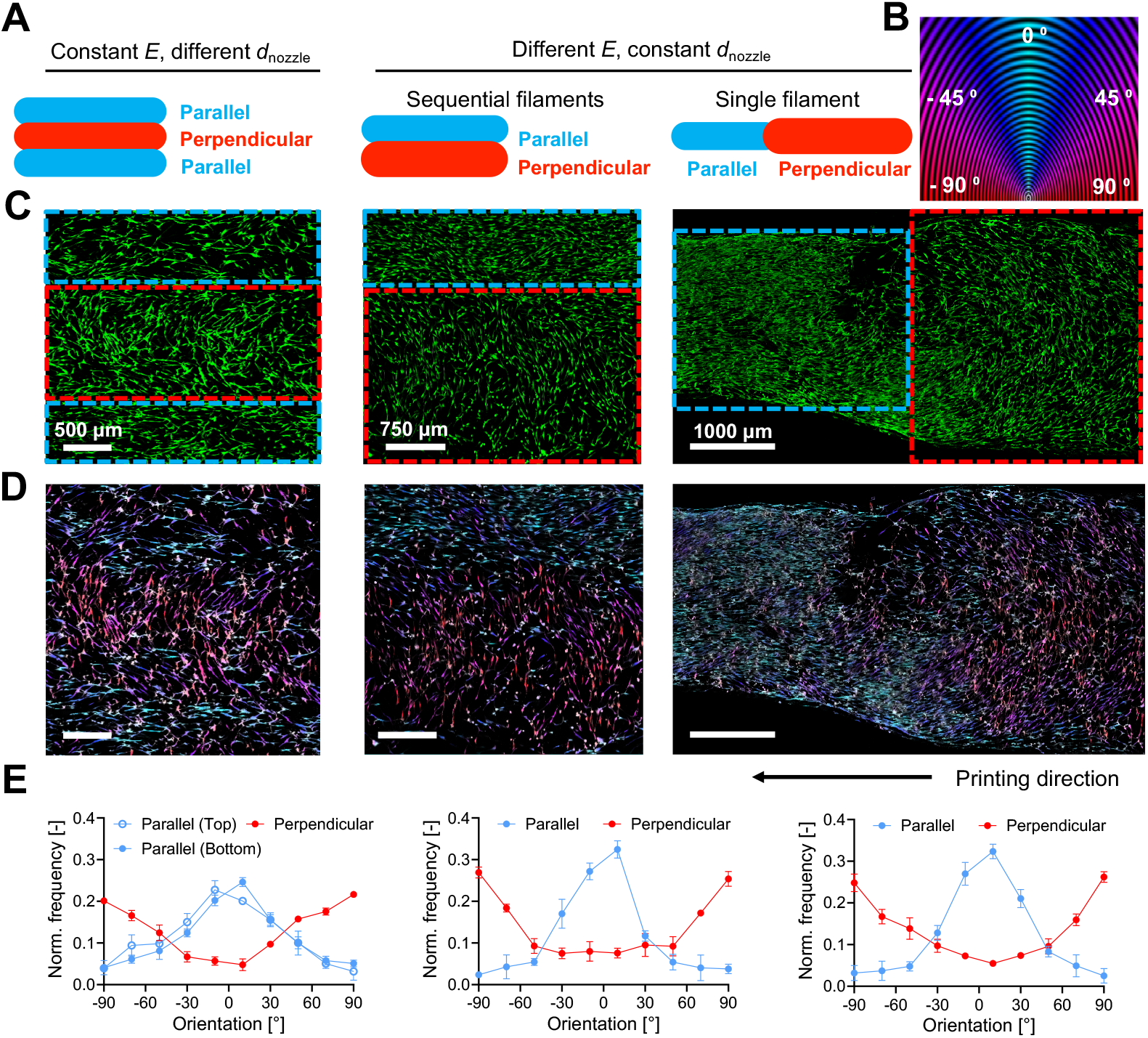
Multidirectional fiber and cell alignment in a single print. (A) Schematic of designs used to achieve multidirectional collagen fiber alignment in a single print, where the blue and red colors refer to regions printed with *V*^*^ values shown to result in a parallel and perpendicular alignment, respectively. (B) Color code used for automated analysis of cell orientation, relative to the printing direction (0^°^). (C) Representative confocal images of CMSCs cultured on top of the collagen prints, visualized through calcein-AM. (D) Representative images of cell morphology with automated coloring based on their orientation using the legend from panel B. (E) Histograms of cell orientation for the different regions of each design, normalized to the total cell count.

## 3. Conclusion

Control over fiber alignment in engineered scaffolds is essential to accurately emulate natural tissue. Here, we demonstrate an extrusion-based 3D printing approach to control collagen fiber orientation in multiple directions by tuning the printing parameters. Importantly, CMSCs cultured on the printed collagen scaffolds were found to orient along the direction of the patterned collagen fibers across multiple days. While here we used plunger-controlled 3D printing, the same principles should be applicable to other types of extrusion-based 3D printing, such as a pressure-controlled setup (65). The key insight is that post-extrusion fluid movement of the ink can induce alignment of embedded fibers. When the printing parameters are selected so that the ink undergoes lateral filament spreading, the fibers can align perpendicular to the printing direction. In contrast, when the printing parameters are selected so that the ink undergoes extensional extensional drawing, the fibers will align parallel to the printing direction. Using conservation of mass principles and automated quantification of fiber alignment, we were able to identity the printing parameters that reliably produce parallel, random, and perpendicular fiber patters. Specifically, we identified *V*^*^ (the speed of ink extrusion from the nozzle relative to the translational speed of the printhead) as being able to directly modify *D*^*^ (the filament diameter normalized to the nozzle diameter), and hence control fiber alignment. From equations 8 and 15, *V*^*^ is related to *D*^*^ as:

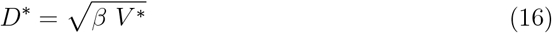

where *β* is a material-specific constant that will likely depend on intrinsic ink properties such as extensional viscosity. For the high concentration collagen ink used in this study, we experimentally determined *β* ≈ 4. Future work to identify the range of ink properties that are compatible with this fiber alignment strategy may allow extension to other types of fibrous inks. For example, our 3D printing alignment strategy could be applied to other fibrous or anisotropic materials, such as cellulose nanofibers, filamentous bacteriophages, or composites of fibrous and amorphous collagen inks (66; 67; 68), which would likely have distinct *β* constants.

The test structures printed here were limited primarily to planar constructs, as this allowed for easy visualization and quantification of fiber alignment. Layer-by-layer extrusion printing has been successfully used to fabricate a wide variety of different 3D constructs with more complex geometric features. As a first step towards this goal, we demonstrated that lateral filament spreading and extensional filament drawing can both be achieved within a support bath to induce successful fiber alignment in the perpendicular and parallel directions, respectively. Future work could build on these proof-of-concept experiments to fabricate structures with more complex 3D shapes that have different fiber alignment patterns in different regions of the tissue. Furthermore, multiplexing this extrusion-based printing strategy with other printing modalities would open up further possibilities to achieve biomimetic structures.

Here, we demonstrate that the patterned fibrous structures were able to induce cellular alignment of the seeded CMSCs, which was maintained over five days. Cell morphology is known to correlate with a number of changes in cell phenotype, and future studies could explore how the fibrous patterns might impact cell proliferation and migration rates, in addition to possible changes in protein expression. Moreover, a great deal of other cell types have been reported to respond to aligned fibrous structures, including endothelial cells, neurons, and myoblasts, to name only a few (69; 70; 71; 72; 73; 74; 75). Thus the 3D printing strategy demonstrated here could be readily extended to a wide range of other cell types. For our studies, cells were seeded onto the surfaces of pre-fabricates constructs; therefore, it remains to be explored if cells could also be directly incorporated into the ink prior to printing to formulate a living bioink. Our collagen ink is already at physiological pH inside the print cartridge; thus, the ink material is cell compatible. On the other hand, the presence of cells may significantly alter the viscoelastic ink properties, which may interfere with fiber alignment. Alternative strategies might include separate printing of an acellular collagen ink and a cellular ink using two print nozzles, as has been previously demonstrated for embedded printing (29; 76).

In summary, this work demonstrates a theoretical and experimental framework to achieve the reproducible fabrication of 3D printed structures with controlled collagen fiber patterns. This technique can be used to achieve different patterns of collagen fiber alignment within different regions of a fabricated construct. These patterns of fiber alignment then induce cell orientation along pre-determined directions. This multidirectional fiber alignment strategy greatly expands the design space for 3D printing of complex, biomimetic tissues.

## 4. Materials and methods

### 4.1. Materials

The collagen ink used was the commercial bovine type I neutralized 35 mg mL^−1^ Lifeink 200 (Advanced Biomatrix). Carbopol ETD2020 powder (Lubrizol) was dissolved by rotating overnight at room temperature at 7 mg mL^−1^ in 100 mL of sterile, ultrapure deionized water (Millipore) with 0.8 mL of 10 M NaOH (Sigma) to balance the pH to 7. Subsequently, the solution was degassed overnight before use.

### 4.2. Rheometry

The mechanical properties of the collagen and Carbopol were evaluated through rheological characterization. Small-amplitude oscillatory shear and rotational measurements were performed on an ARG2 rheometer (TA Instruments) equipped with a Peltier plate and a solvent trap to prevent evaporation, using a parallel-plate geometry with a diameter of 8 mm. The absence of wall slip was confirmed through measurements at a different measuring gap having excellent agreement with each other. Collagen gelation was examined through time-sweep measurements at an angular frequency of 1 rad s^−1^ and a shear strain amplitude of 1% at a temperature between 4^°^C and 37^°^C. Frequency-sweep measurements were conducted after the storage modulus (*G*^′^) had reached a plateau at a shear strain amplitude of 1%, in the linear viscoelastic regime, between an angular frequency of 0.1 and 100 rad s^−1^. Step-shear measurements were conducted with an alternating shear strain amplitude of 0.1% and 300% using a parallel-plate geometry with a diameter of 8 mm and a measuring gap of 900 *µ*m and at an angular frequency of 6 rad s^−1^. For Carbopol each shear step lasted 30 s and the measurements took place at room temperature, while for the collagen ink the low-shear step lasted 300 s and the high-shear step 40 s and the measurements were conducted at 4^°^. Three replicates using a fresh sample were conducted for all measurements.

### 4.3. 3D Printing

3D printing was performed using a custom-built dual-extruder bioprinter modified from a MakerGear M2 Rev E plastic 3D printer, as previously described [8, 58, 59]. Briefly, the thermoplastic extruder of the printer was removed and replaced with a mount designed to hold two Replistruder 4 syringe pumps. Additionally, the control board was replaced with a Duet 2 WiFi board with RepRapFirmware. The syringe was wrapped with Gel Finger Ice Pack (stored at -20 °C) to retain the collagen at 4^°^C, as suggested by the manufacturer, to keep the self-assembly kinetics constant during printing. The printing was performed with nozzles having a diameter of 1.55 mm (14G), 0.84 mm (18G), 0.42 mm (22G), and 0.16 mm (30G) and the plunger distance varied from *E* = 0.003 mm to *E* = 0.04 mm. The printing speed (F) was kept constant at 99920 mm min^−1^, the step distance (Δ*x* = 0.5 mm) and syringe diameter (D = 7.29 mm) for all samples. The entire printing took place in air at room temperature. Acellular prints were printed as a single 2 cm long filament while the diameter varied depending on the printing parameters ranging from (1 mm to 4.8 mm). The Gcode used were manually written in a text file. For prints that were subsequently seeded with cells, the printing process was carried out in a sterile tissue culture cabinet and each sample consisted of three sequentially printed filaments with a length of 2 cm to increase seeding area. For perpendicular alignment, the sample was printed using a *d*_nozzle_ of 0.42 mm (22G) with *E* = 0.007 mm and for parallel alignment using a *d*_nozzle_ of 1.55 mm (14G) with *E* = 0.007 mm.

### 4.4. Cell culture

Corneal mesenchymal stromal cells (CMSCs) were obtained from a human donor cornea (Lions Eye Institute for Transplant and Research) following established protocols (77). The donor, aged between 30 and 35 years, had no known history of herpes simplex virus, varicella zoster virus, human immunodeficiency virus, or hepatitis. The time from death to preservation was less than 7 days, and the cornea was stored in Optisol corneal storage medium. Cells were cultured in a growth medium containing 500 mL MEM-Alpha (Corning), 50 mL fetal bovine serum (Gibco), 5 mL GlutaMax (Gibco), 5 mL non-essential amino acids (Gibco), and 5 mL antibiotic-antimycotic (Gibco). The growth medium was refreshed every other day, and CMSCs were passaged once they reached 80% confluency. The cells were used in experiments between passages 6 and 10. The cells were seeded on the bottom side of the print as it is more flat than the top part of the print. To achieve this, the prints were flipped in PBS solution followed by pipetting out the PBS solution. Then, CMSCs were trypsinized, counted, pelleted, re-suspended in media, and seeded at an initial cell density of 10^4^ cells cm^−2^.

### 4.5. In-vitro Characterization

To assess the viability of the CMSCs, a Live/Dead assay was conducted by staining cells with calcein AM and ethidium homodimer-1 (Life Technologies), following the manufacturer’s instructions. Imaging for live/dead analysis was performed using a STELLARIS 5 confocal microscope (Leica) with a 10X air objective. At least 5 images were taken in different areas of each print, and image analysis was performed using FIJI (ImageJ2, Version 2.3.0/1.53f). Cell viability was calculated as the number of live cells (calcein AM-positive) divided by the total number of cells.

For immunofluorescence imaging, gels with seeded CMSCs were fixed with 4 % paraformaldehyde (PFA) in PBS for 20 minutes at room temperature (RT) followed by three washes (10 minutes each). Samples were permeabilized for 45 minutes at RT with 0.25 % Triton X-100 (Sigma Aldrich) in PBS (PBST) and then blocked for 2 hours at RT using blocking solution containing PBS with 5 wt % bovine serum albumin (BSA, Roche), 5 % goat serum (Gibco), and 0.5 % Triton X-100. Antibody dilutions were prepared in PBS with 2.5 wt % BSA, 2.5 % goat serum, and 0.5 % Triton X-100. Each sample was first treated with the appropriate primary antibody overnight at 4^°^C: for CMSCs, rabbit anti-ALDH-3A1 (Abcam, ab76976, 1:200 dilution). The following day, the samples were washed three times with PBST (20 minutes each). The appropriate dilution of secondary antibodies was prepared in antibody dilution solution: 4’,6’-diamidino-2-phenylindole (DAPI, Molecular Probes, 1:900 dilution) and phalloidin-tetramethyl rhodamine B isothiocyanate peptide from Amanita phalloides (Phalloidin-TRITC, Sigma Aldrich, 1:1000 dilution), Alexa Fluor 488 Goat anti-Rabbit Secondary Antibody, 1:500 dilution) and then added to the samples incubated overnight at 4^°^C. Finally, samples were washed with PBST three times and imaged with a STELLARIS 5 confocal microscope (Leica) using 40x oil and 10x air objectives. Tile scans were taken at 512 x 512 image format.

### 4.6. Microscopy and image analysis

Collagen fibers in the acellular prints were imaged with a STELLARIS 5 confocal microscope (Leica) using 40x oil objective. Tile scans were taken at 512 x 512 image format for the entire print. Confocal images were then analyzed to characterize collagen fiber alignment qualitatively and quantitatively. For qualitative analysis, CT-FIRE for Individual Fiber Extraction software was used to enable automated tracing of the collagen fibers by artificially coloring individual fibers to aid in visual assessment of the overall collagen fiber alignment in the sample (78). For quantitative analysis, confocal images were analyzed using theOrientation J plugin in FIJI (Image J) software. The fraction of fibers corresponding to each orientation was obtained as a function of orientation angle (in degrees,^°^). Using this, histograms of normalized frequency of fibers relative to the total number of fibers versus orientation angle were plotted with bins of 20^°^. Three independent printing trials were analyzed for each experimental condition and plotted as mean ± standard deviation. Cellular alignment of the seeded CMSCs on the gels was analyzed with CellProfiler, an open-source software using confocal images of the cells visualized with calcein-AM (live) stain.

### 4.7. Statistical analysis

Statistical analyses were performed using GraphPad Prism (Version 10). Details of the sample sizes and statistical tests conducted for each figure are included within the figure captions. In all cases, statistical differences are denoted as follows: not significant (ns *p* > 0.05),^*^*p* < 0.05^**^*p* < 0.01,^***^ *p* < 0.001,^****^ *p* < 0.0001).

## Acknowledgments

The authors thank Betty Cai and Dr. David Kilian for the helpful discussions. The authors acknowledge funding support from the Stanford Graduate Fellowship in Science and Engineering (D.S.), the Swiss National Science Foundation including P500PN210723 (F.C.), the National Institutes of Health including R01-EY035697 (S.C.H., D.M.), F31-EY034785 (L.G.B), and K01-EB033870 (V.M.D.), and the ARCS Foundation Scholarship (L.G.B.).

