## Supplemental Information for "Multidirectional alignment of collagen fibers to guide cell orientation in 3D-printed tissue"

**Diya Singhal[1,‡], Fotis Christakopoulos[2,‡], Lucia G. Brunel[1], Suraj Borkar[1], Vanessa M. Doulames[2], David Myung[3,4], Gerald G. Fuller[1] and Sarah C. Heilshorn[2,*]**

[1] Department of Chemical Engineering, Stanford University, Stanford, CA, USA
[2] Department of Materials Science and Engineering, Stanford University, Stanford, CA, USA
[3] Department of Ophthalmology, Byers Eye Institute, Stanford University School of Medicine, Palo Alto, CA, USA
[4] VA Palo Alto Healthcare System, Palo Alto, CA, USA
‡ These authors contributed equally to this work

**List of Supplemental Information**:

**Figure S1**. Characterization of collagen ink.
**Figure S2**. Schematic of the extrusion-based printing setup.
**Figure S3**. Representative fiber traces.
**Figure S4**. Filaments printed with a nozzle diameter of 0.42 mm
**Figure S5**. Histograms of fiber alignment keeping $E$ constant.
**Figure S6**. Multi-filament prints with aligned collagen fibers and cells.
**Figure S7**. Viability of CMSCs cultured on patterned collagen.
**Table S1**. Printing parameters for the prints used in the study.
**Derivation for velocity ratio, $V^*$.**

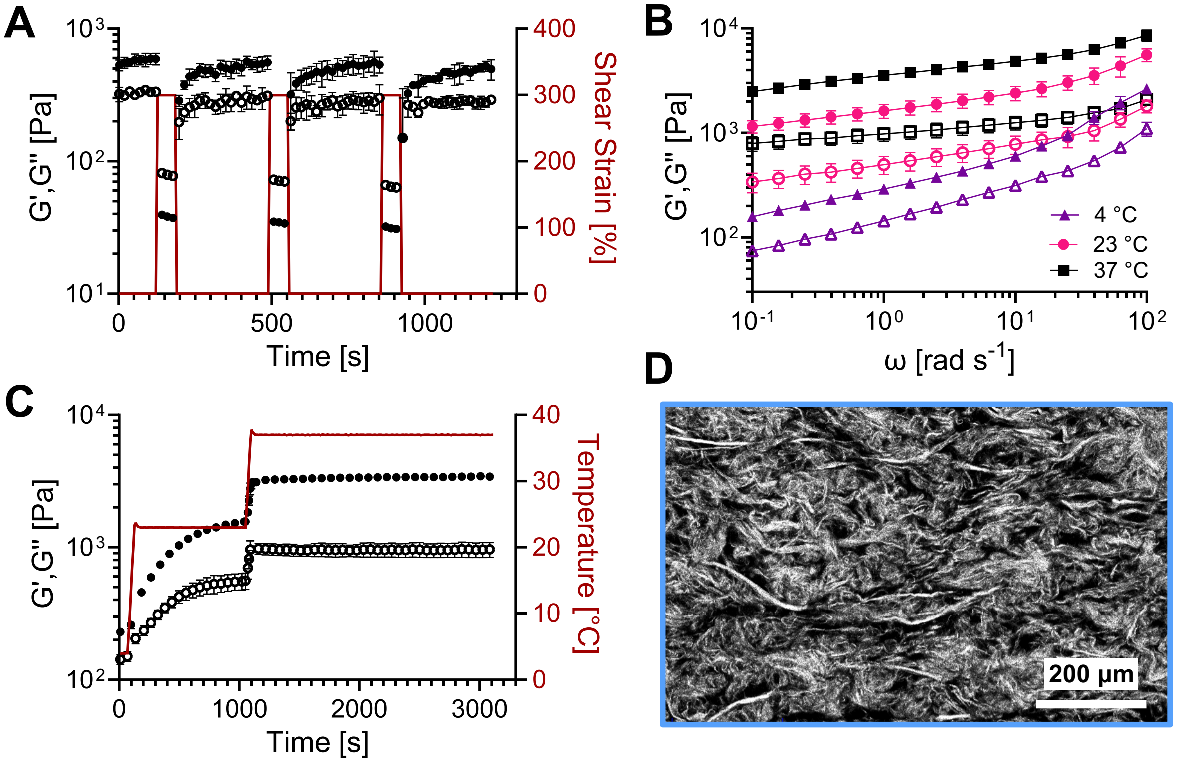


Figure S1: Characterization of collagen ink. For all plots, filled symbols correspond to storage moduli, $G'$, and open symbols correspond to loss moduli, $G''$ (A) Shear rheology at 4 °C during alternating application of high or low shear strain (red line) demonstrates shear-thinning and self-healing behavior. (B) Frequency-sweep of the collagen ink at 4 °C (purple), 23 °C (pink), and 37 °C (black). (C) Shear rheology of the collagen ink as temperature increases from 4 to 37 °C. (D) Representative confocal reflectance image of the collagen fibers post-printing after incubation at 37 °C for 15 minutes.

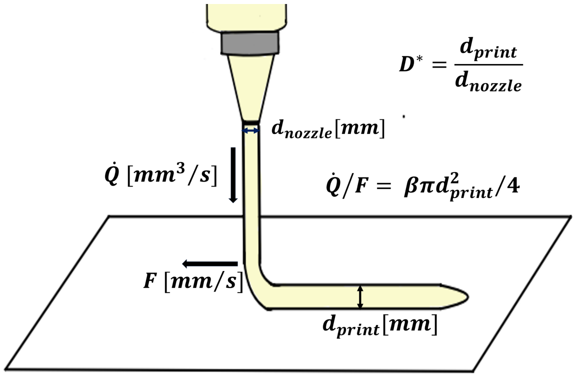


Figure S2: Schematic of the extrusion-based printing setup. Key printing parameters and equations relevant for the calculation of $D^*$ are shown; nozzle diameter ($d_{\text{nozzle}}$, mm), translational speed of the printhead ($F$, mm min$^{-1}$), volumetric flow rate ($\dot{Q}$, mm$^3$ $s^{-1}$), and diameter of the printed filament ($d_{\text{print}}$).

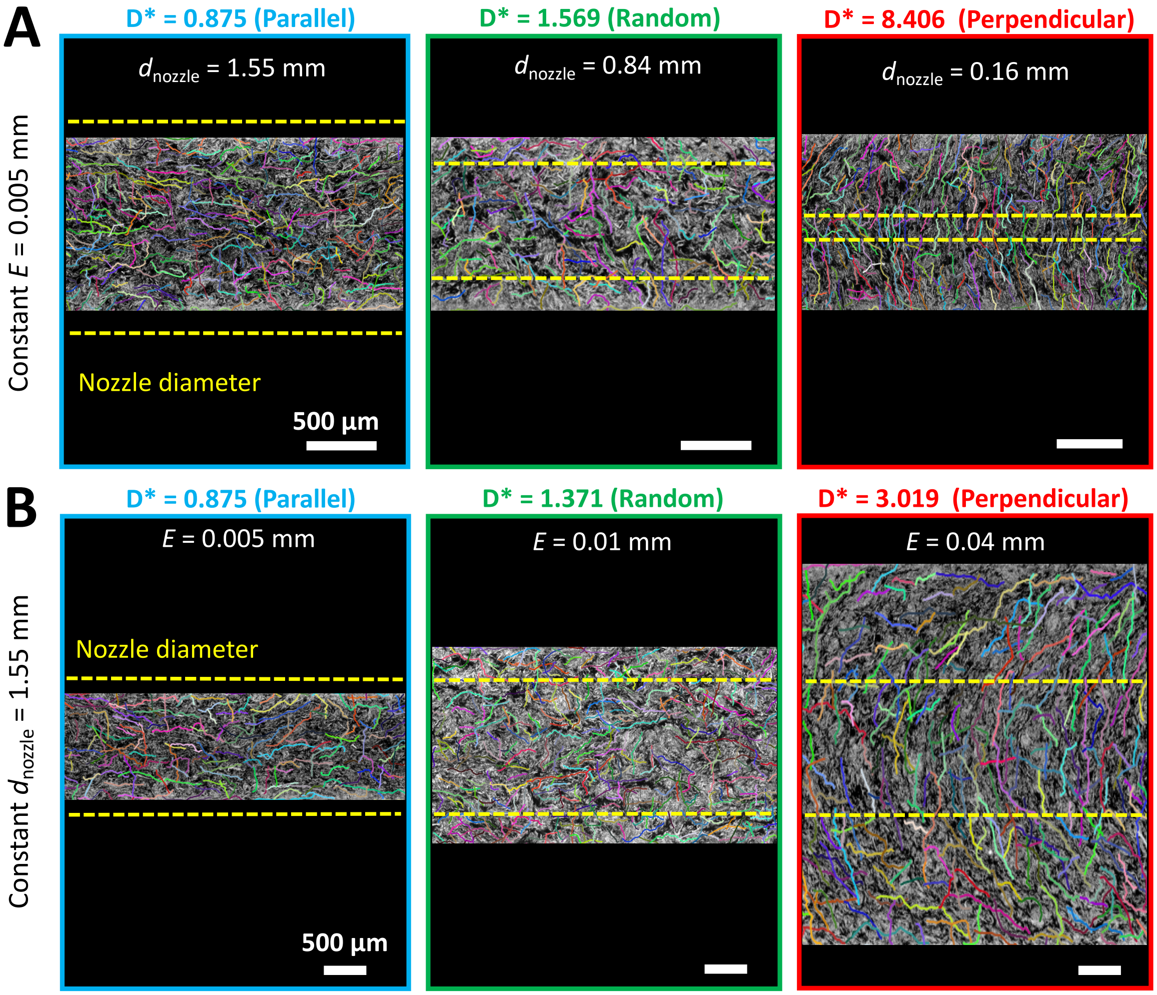


Figure S3: Representative fiber traces. Images are repeated from figure 2 with individual collagen fibers traced and artificially colored through the automated fiber tracing algorithm, CT-FIRE for Individual Fiber Extraction. (A) Representative confocal reflectance images of collagen filaments printed with the same plunger displacement per step ($E = 0.005$ mm) and different nozzle diameters ($d_{\text{nozzle}}$). (B) Collagen filaments printed with the same nozzle ($d_{\text{nozzle}} = 1.55$ mm) and different plunger displacement per step ($E$). Yellow dotted lines denote the nozzle diameter for easy comparison to the printed filament width. The experimentally determined diameter ratio, $D^*$, (*i.e.* the width of the printed filament, $d_{\text{print}}$, normalized by the diameter of the printing nozzle, $d_{\text{nozzle}}$) is written above each micrograph along with a qualitative description of the observed collagen fiber orientation. Scalebars correspond to 500 $\mu$m.

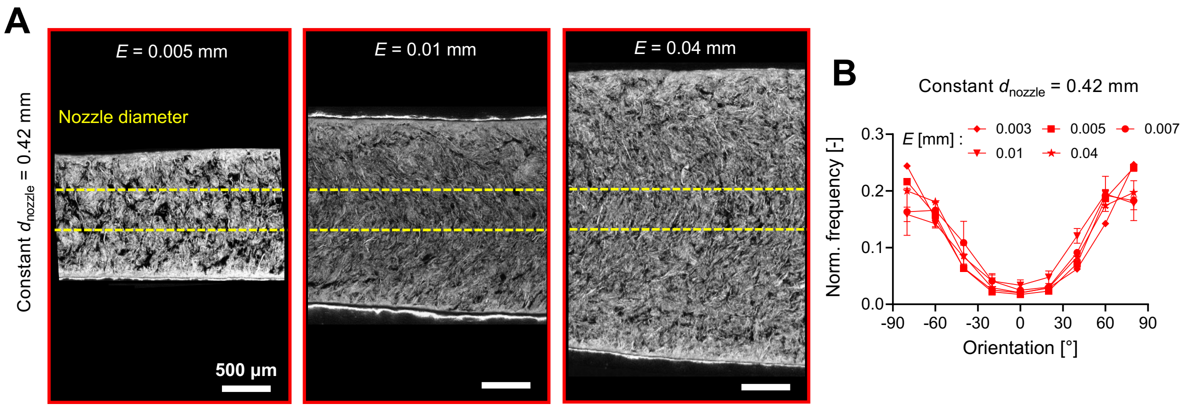


Figure S4: Filaments printed with a nozzle diameter of 0.42 mm. (A) Representative confocal reflectance images of collagen filaments printed with the same nozzle diameter ($d_{\text{nozzle}} = 0.42\ mm$) and different plunger displacement per step, $E$. Yellow dotted lines denote the nozzle diameter for easy comparison to the printed filament width. (B) Histogram of fiber alignment with keeping $d_{\text{nozzle}}$ constant (0.42 mm) and changing $E$, showing the normalized frequency of fiber alignment relative to the total number of fibers versus fiber orientation, where the printing direction is 0 °. (B) Data are mean ± standard deviation, N=3 independent printing trials.

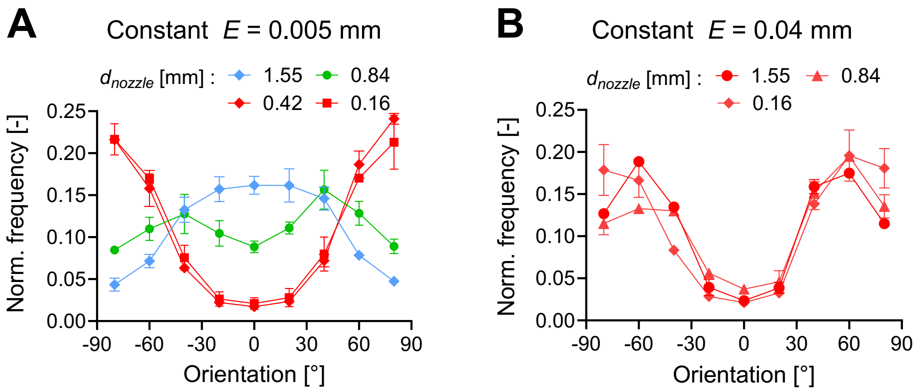


Figure S5: Histograms of fiber alignment with keeping $E$ constant. (A) For $E$ equal to 0.005 mm and (B) for $E$ equal to 0.04 mm and changing $d_{\text{nozzle}}$, show the normalized frequency of fiber alignment relative to the total number of fibers versus fiber orientation, where the printing direction is 0 °. (A-B) Data are mean ± standard deviation, N=3 independent printing trials.

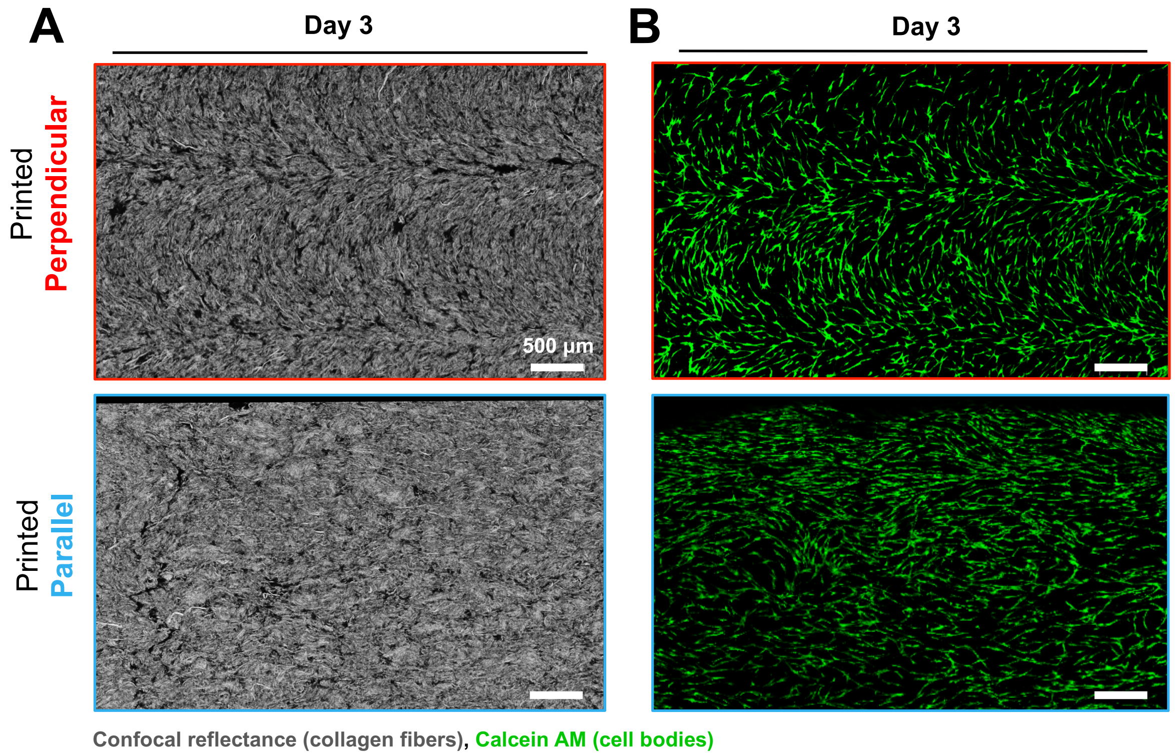


Figure S6: Multi-filament prints with aligned collagen fibers and cells. Each print consists of multiple filaments of same alignment pattern to increase seeding area. (A) Representative confocal reflectance images of collagen fibers aligned in the perpendicular (top) and parallel (bottom) directions. (B) Representative confocal images on day 3 of calcein-AM stained CMSCs cultured on the collagen prints with underlying collagen fibers aligned in the perpendicular (top) and parallel (bottom) directions.

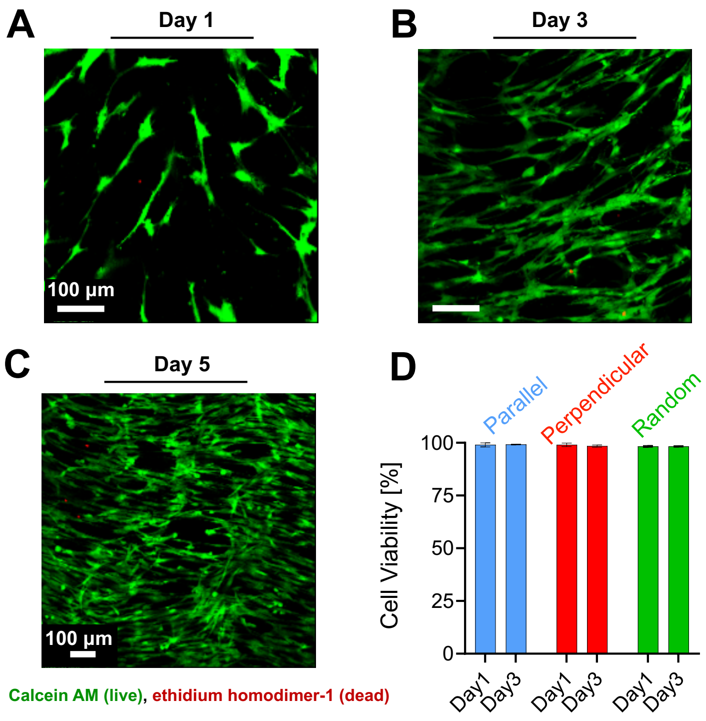


Figure S7: Viability of CMSCs cultured on collagen scaffolds. (A-C) Representative confocal images from Live/Dead viability assay of CMSCs cultured on collagen with perpendicular fiber alignment stained for on day 1 (A), day 3 (B), and day 5 (C). (D) Cell viability for CMSCs cultured on collagen scaffolds with different fiber alignment patters on days 1 and 3. High cell confluence at day 5 prevented automated identification of cell boundaries

<table>
<tr><th>Figure / description</th><th>$d_{\text{nozzle}}$ [mm]</th><th>$E$ [mm]</th><th>$D^*$ [-]</th><th>$V^*$ [-]</th><th>Print geometry</th></tr>
<tr><td>Figure 1(F)</td><td>1.55</td><td>0.005</td><td>0.874</td><td>0.220</td><td>1 filament</td></tr>
<tr><td>Figure 5(C),<br>print in support bath</td><td>1.55</td><td>0.007</td><td>1.04</td><td>0.303</td><td>1 filament</td></tr>
<tr><td>Figure 5(D),<br>print in support bath</td><td>0.42</td><td>0.007</td><td>4.25</td><td>4.185</td><td>1 filament</td></tr>
<tr><td>Figure 6(A)</td><td>0.42</td><td>0.007</td><td>4.25</td><td>4.185</td><td>3 sequential filaments</td></tr>
<tr><td>Figure 6(A)</td><td>1.55</td><td>0.007</td><td>1.04</td><td>0.303</td><td>3 sequential filaments</td></tr>
<tr><td rowspan="2">Figure 7(C),<br>left print,<br>constant $E$, different $d_{\text{nozzle}}$</td><td>1.55<br>(top, bottom)</td><td rowspan="2">0.007</td><td>1.04<br>(top, bottom)</td><td>0.303<br>(top, bottom)</td><td rowspan="2">3 sequential filaments</td></tr>
<tr><td>0.42<br>(middle)</td><td>4.25<br>(middle)</td><td>4.185<br>(middle)</td></tr>
<tr><td rowspan="2">Figure 7(C),<br>middle print,<br>different $E$, constant $d_{\text{nozzle}}$</td><td rowspan="2">1.55</td><td>0.007<br>(top)</td><td>1.04<br>(top)</td><td>0.303<br>(top)</td><td rowspan="2">2 sequential filaments</td></tr>
<tr><td>0.040<br>(bottom)</td><td>3.02<br>(bottom)</td><td>1.780<br>(bottom)</td></tr>
<tr><td rowspan="2">Figure 7(C),<br>right print,<br>different $E$, constant $d_{\text{nozzle}}$</td><td rowspan="2">1.55</td><td>0.007<br>(left)</td><td>1.04<br>(left)</td><td>0.303<br>(left)</td><td rowspan="2">1 filament</td></tr>
<tr><td>0.040<br>(right)</td><td>3.02<br>(right)</td><td>1.780<br>(right)</td></tr>
</table>

Table S1: Printing parameters for the prints used in the study. Each filament in the print has a length of 2 cm. Total print width is obtained by multiplying the number of sequential filaments with print width of single filament. The print width of single filament can be calculated as a product of $d_{\text{nozzle}}$ and $D^*$.

### Derivation for Velocity Ratio, (V*)

We defined the normalized velocity, $V^*$:

$$V^* = \frac{C}{F} \tag{S1}$$

where $C$ corresponds to the speed of the ink extrusion from the nozzle and $F$ to the translational speed of the printhead. For our extrusion-based, piston-controlled, 3D printer, the volumetric flow rate inside the syringe ($\dot{Q}_{\text{syringe}}$) is defined as:

$$\dot{Q}_{\text{syringe}} = \frac{A_{\text{syringe}} E}{\frac{\Delta x}{F}} \tag{S2}$$

which, due to mass conservation, is equal to the volumetric flow rate at the end of the nozzle ($\dot{Q}_{\text{nozzle}}$):

$$\dot{Q}_{\text{nozzle}} = \dot{Q}_{\text{syringe}} \tag{S3}$$

The volumetric flow rate at the end of the nozzle can be defined as:

$$\dot{Q}_{\text{nozzle}} = C\frac{\pi d_{\text{nozzle}}^2}{4} \tag{S4}$$

Using supplemental equation S3, we find that:

$$\frac{A_{\text{syringe}}E}{\frac{\Delta x}{F}} = C\frac{\pi d_{\text{nozzle}}^2}{4} \tag{S5}$$

Thus, we can derive velocity ratio ($V^*$) for our extrusion based 3D printer setup as:

$$V^* = \frac{C}{F} = \frac{4A_{\text{syringe}}E}{\pi d_{\text{nozzle}}^2 \Delta x} \tag{S6}$$
